# GLESKit: an MCP-based system for gene-level biomedical investigation and evidence synthesis

**DOI:** 10.64898/2026.10.25.761689

**Authors:** Athos Moraes, Pedro G. Ferreira

## Abstract

Biomedical investigation at the gene level requires researchers to combine expression data, functional annotations, pathways, interaction networks, genetic associations, drug evidence and literature. Accessing these resources involves databases, analysis tools and scripts that require bioinformatics expertise, complicating the production of clear, reviewable reports. We present GLESKit, a gene centred system that connects these steps through the Model Context Protocol and retains data, analyses, sources and reports within a study workspace. We evaluated report production with 15 author curated GeneReportBench tasks and question answering with a gene level subset of BixBench, a reference biology benchmark for agentic systems. GLESKit completed all 15 reports with a mean committee score of 0.904, although evidence quality varied, and earned positive credit on 35 of 39 scored human BixBench questions with a mean of 0.438. In paired tests, configurations with skills scored 0.145 higher across 20 questions and 0.085 higher across five reports.

---

Understanding how genes shape cellular states, disease mechanisms and therapeutic responses is central to biomedical research. Foundation models trained on single-cell transcriptomes support context-aware gene- and cell-level predictions, with model scaling and quantization shaping their computational requirements [1]. Gene-level investigation provides a common scale at which transcriptomic measurements can be related to protein function, molecular pathways, interaction networks, genetic associations and drug evidence [2]. The increasing availability of high-throughput assays and public biomedical resources has expanded what researchers can learn from a single gene, a set of genes or a differential-expression analysis, while making interpretation increasingly dependent on computational analysis. Converting these heterogeneous observations into a defensible biomedical account requires researchers to connect analytical steps, preserve the biological context and provenance of intermediate outputs, and qualify the evidence supporting each conclusion.

This interpretive burden is especially demanding when researchers bring deep biomedical knowledge but limited training in the software and data conventions of bioinformatics. Clinicians, pathologists and experimental biologists may be well positioned to judge whether a result is plausible or useful, while still depending on computational specialists to configure analyses, reconcile identifiers, query multiple services and prepare figures. Platforms that make computational biology more accessible have consequently treated usability, reproducibility and reuse as central design requirements [3, 4]. Natural-language interaction could help bridge domain and computational expertise by allowing researchers to formulate questions in familiar terms. The scientific value of natural-language interaction depends on preserving the operations, analytical context and source material required for expert inspection.

Several generations of computational systems address parts of this workflow. Taverna [5], GenePattern [3] and Galaxy [4] support the composition and execution of multi-step bioinformatics analyses through reusable workflows, graphical access and shared computational infrastructure. GenomeSpace [2] demonstrated interoperation among genomic-analysis tools, whereas the GenePattern Notebook Environment [6] and BioJupies [7] connected computation with narrative, figures and report-like outputs. Tool-using language models contribute to a complementary mode of interaction. ReAct [8] interleaves reasoning with external action, Toolformer [9] showed that models can learn to invoke API-like tools, and GeneGPT [10] connected a language model to NCBI services for genomics question answering. Benchmarking of agentic viral-sequence retrieval [11] further showed that a deterministic, reproducible programmatic interface improved accuracy and response stability across multiple agent systems, illustrating how data-access design can constrain scientific-agent reliability. Biomni [12] extends this direction through a general-purpose biomedical action environment that combines resource retrieval, code-based execution and adaptive planning across databases, software and experimental protocols. In a different scientific setting, an agentic X-ray scientist [13] uses tools exposed through the Model Context Protocol and persistent experimental state to conduct closed-loop instrument operations with existing reasoning models. Other systems integrate scientific tools, persistent workspaces, hypothesis generation or more autonomous experimental and computational research [14, 15, 16, 17, 18]. Collectively, these systems position tools, skills, execution environments and persistent records as core components of scientific-agent capability. Human-in-the-loop (HITL) gene-level analysis adds a specific integration challenge: biomedical operations must remain accessible across models and conversational environments, accommodate changing tools and data modalities, and connect analyses, evidence and reports throughout an evolving investigation. This integration is particularly relevant for researchers with biomedical expertise who rely on computational support for bioinformatics workflows.

Here we present GLESKit (Gene-Level Evidence Synthesis Toolkit), an integrated environment designed for HITL gene-level biomedical investigation. The system uses genes as its organizing core, linking gene-expression results, gene and protein annotations, genetic associations and drug–target evidence through normalized gene identifiers while preserving each result’s source and evidentiary type. A modular tool layer connects established bioinformatics resources and analytical methods for retrieval, analysis, visualization and reporting behind a common interface (Extended Data Table 5), a design that focuses the LLM on coordinating and synthesizing tool-derived evidence instead of relying on its internal knowledge alone. GLESKit makes that interface available through the Model Context Protocol (MCP) [19], separating the biomedical operations from any one model or conversational client and permitting their use through a web application served on the user’s own computer and accessed in a browser, or through another compatible client. Provider adapters allow new resources and analytical functions to be added while preserving the client-facing interface. The optional web interface adds persistent study sessions, configurable report templates and stored intermediate or final research outputs so that analyses and evidence can contribute to an inspectable initial synthesis. In intended use, a primary conversational agent maintains task context and invokes tools while the researcher directs the investigation and reviews its interpretations. We report two benchmark evaluations and paired skills ablations. GeneReportBench, our structured biomedical report benchmark, assesses completion, structure and evidentiary support in extended report workflows. A scope-filtered subset of BixBench [20] evaluates focused computational-biology answers and execution outcomes. Finally, paired ablations evaluate matched BixBench questions and GeneReportBench reports with and without skills. Together, these automated evaluations characterize report production, focused answer quality, skills and the system’s current operating scope.

## Results

### GLESKit integrates biomedical analysis and evidence synthesis around genes

GLESKit provides a information unit and a modular system for investigations that evolve across data, analysis and evidence synthesis. A study can begin with raw RNA-sequencing counts, a differential-expression table, one or more genes, a biomedical question or a hypothesis. From these entry points, the system can resolve gene and protein identities, inspect expression and functional context, perform gene-set enrichment or differential-expression analyses, examine pathways and interaction networks, retrieve disease and drug associations, gather literature, and generate figures or produce a structured report (Fig. 1a,b and Extended Data Table 5). The conversational agent coordinates these operations, while the researcher can redirect the analysis, request additional evidence and decide which outputs contribute to the final account.

**Fig. 1.**
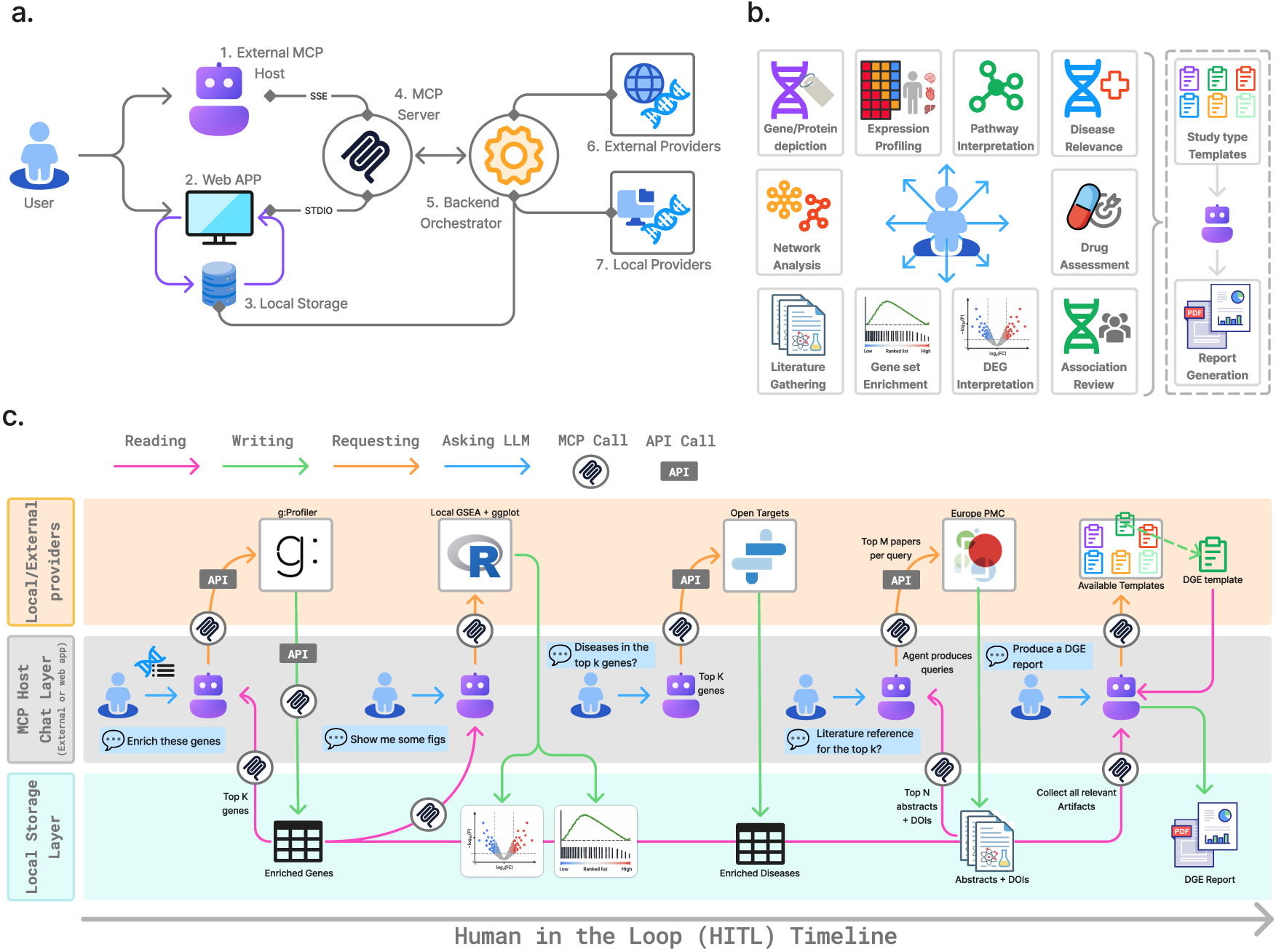
Scope, access and intended HITL workflow in GLESKit. **a**, A researcher can access the same biomedical MCP service through a compatible external host or the locally served web application, opened in a browser to access the workspace and its local storage. The backend orchestrator connects MCP-exposed tools to local and external providers. **b,** Representative gene-level capabilities are implemented as tool-backed biomedical operations that can be assembled into study-specific reports. **c,** The intended HITL workflow progresses from differential-expression analysis through gene-set enrichment, figure generation, disease associations and literature review to a template-guided report, with intermediate outputs retained throughout the investigation. The benchmark runs reported here were fully automated.

Genes provide a normalization anchor across these operations. Transcriptomic measurements, protein annotations, genetic associations and pharmacogenomic evidence retain their distinct evidentiary types, while normalized gene identifiers relate their outputs within the same investigation. This entity model connects providers and packaged analyses while preserving the source and type of each result. The current implementation concentrates on transcriptomic and gene-level questions about human biology. Effective extension to further modalities or organisms depends on appropriate resources, identifier policies and validation.

The biomedical operations are exposed through MCP as a shared client-facing layer (Fig. 1a). A compatible external host can invoke the same operations, while the web application adds persistent study sessions that connect the conversation with uploaded materials, notes, analyses, evidence, figures and reports. We call these stored intermediate or final research outputs artifacts, with the operational definition provided in Methods. Their association with a study session allows a researcher to inspect and reuse materials as the investigation develops and to assemble them into an initial report for further review (Fig. 1c).

### Complementary benchmarks evaluate focused answers and structured biomedical reports

The evaluation addresses two complementary task types. GeneReportBench asks whether an extended investigation can produce a structured report and preserve materials that make it reviewable. Its 15 tasks span three report types: five biomarker-prioritization, five mechanism-interpretation and five disease-context reports. BixBench [20] measures performance on focused computational-biology questions. The prespecified RNA-centred and gene-level scope contains 67 questions from 20 cases spanning three species: 41 concerning *Homo sapiens*, 13 concerning *Mus musculus* and 13 concerning *Pseudomonas aeruginosa*. The evaluation sequence and reuse of benchmark runs are summarized in Extended Data Fig. 6.

GeneReportBench evaluates extended synthesis and its supporting record, whereas BixBench evaluates focused answer quality and execution outcomes. Committee scores are benchmark-specific weighted means of normalized model-based specialist ratings, with components and weights defined in Methods. Representative cases make these distinct evaluation targets concrete (Fig. 2). In BixBench task bix-36-q5, three completed GLESKit MCP calls produced a quantitative answer that matched the benchmark target and received the maximum normalized committee score of 1.000. In GeneReportBench task repo-6, 22 total tool calls across 12 distinct tools produced an inflammatory-bowel-disease report with 12 linked artifacts. All four report specialists assigned the highest level, yielding the maximum score of 1.000. Both cases retain the connection between the benchmark task, the agent’s execution and the specialist assessment.

**Fig. 2.**
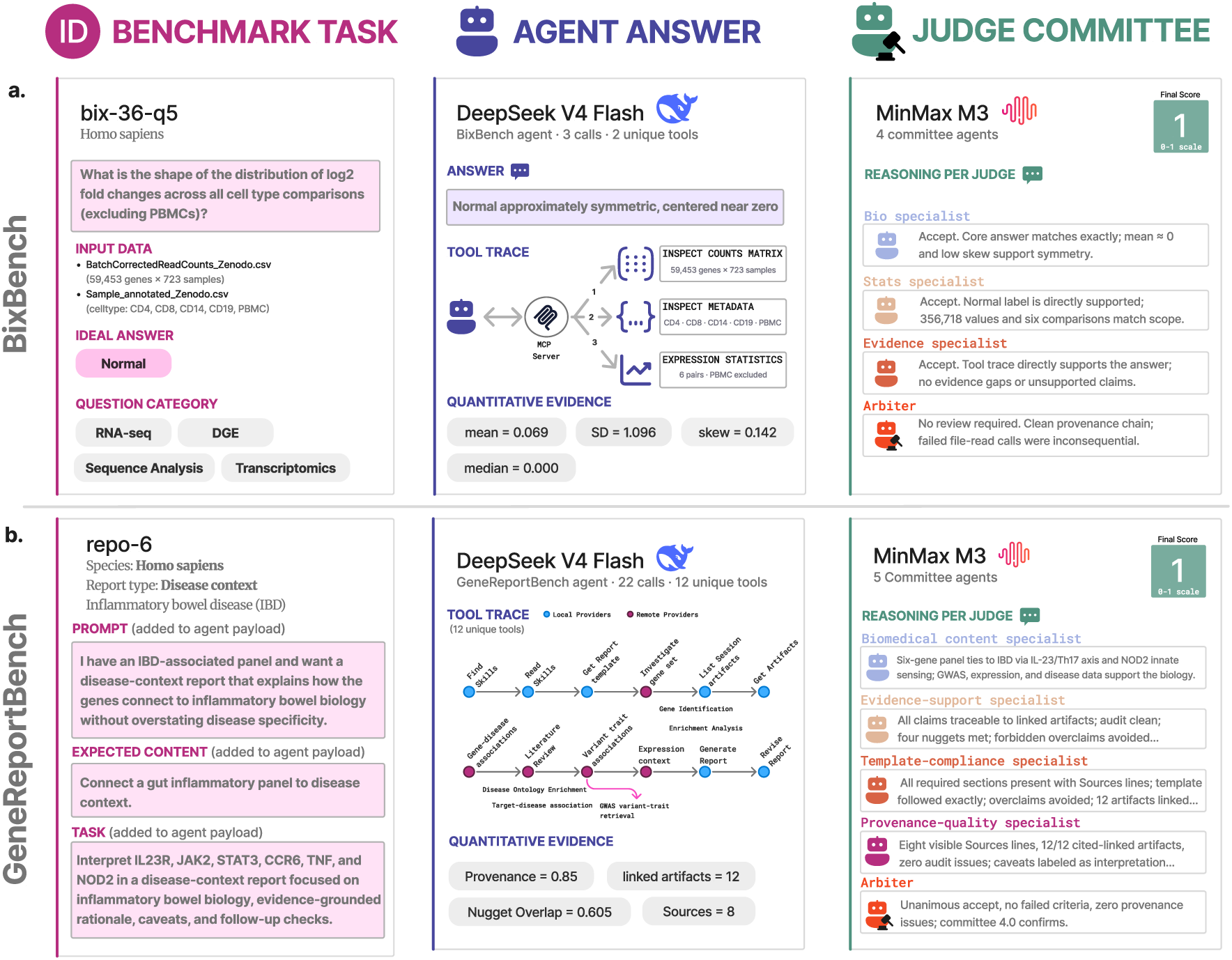
Representative benchmark executions for focused answers and extended reports. **a**, A BixBench question about human biology progresses from the benchmark task through three completed GLESKit MCP calls to a quantitative answer assessed by biology, statistics and evidence specialists and a qualitative arbiter. **b,** A GeneReportBench disease-context task progresses through 22 total tool calls to a saved report assessed for biomedical content, evidence support, provenance quality and template compliance. Both examples received the maximum normalized committee score of 1.000. Panel **b** shows the nugget overlap and provenance values supplied to all four specialists. Committee scores used the four categorical ratings, with no separate weighting for these values (Methods).

Both benchmarks combine recorded workflow measures with benchmark-specific specialist assessment, with the complete selection and scoring protocols described in Methods. Separate paired evaluations examine 20 BixBench questions and five GeneReportBench reports with and without skills, recording answer quality and tool use within each task set.

Outcome classes and analysis denominators are defined in Methods and summarized in Extended Data Table 4.

### All GeneReportBench tasks produced reports with high committee scores, while evidence ratings varied

All 15 GeneReportBench runs completed and produced a saved report with a valid committee score (weighted mean of four normalized model-based specialist ratings). The mean normalized score was 0.904 (95% confidence interval (CI), 0.851–0.958), with an s.d. of 0.097, a median of 0.925 and a range of 0.688–1.000. The mean normalized scores for the three report types were 0.818 for biomarker prioritization, 0.963 for disease context and 0.933 for mechanism interpretation (Fig. 3a).

The specialist assessments resolve the components of the high aggregate score (Fig. 3b). Template compliance was rated Accept for all 15 reports, and biomedical content was Accept for 14 and Mostly Supported for one. Evidence support was less uniform, with five Accept, eight Mostly Supported, one Partial and one Weak rating. Provenance was Accept for nine reports, Mostly Supported for two and Partial for four. The ordered categories and their criteria are defined in Methods. These results reveal a clear gap between consistent structural completion and the variable strength of evidence and provenance. Complete quality outcomes and workflow measures for each task are provided in Extended Data Tables 1 and 2, respectively, with the corresponding records supplied in the Source Data file.

**Fig. 3.**
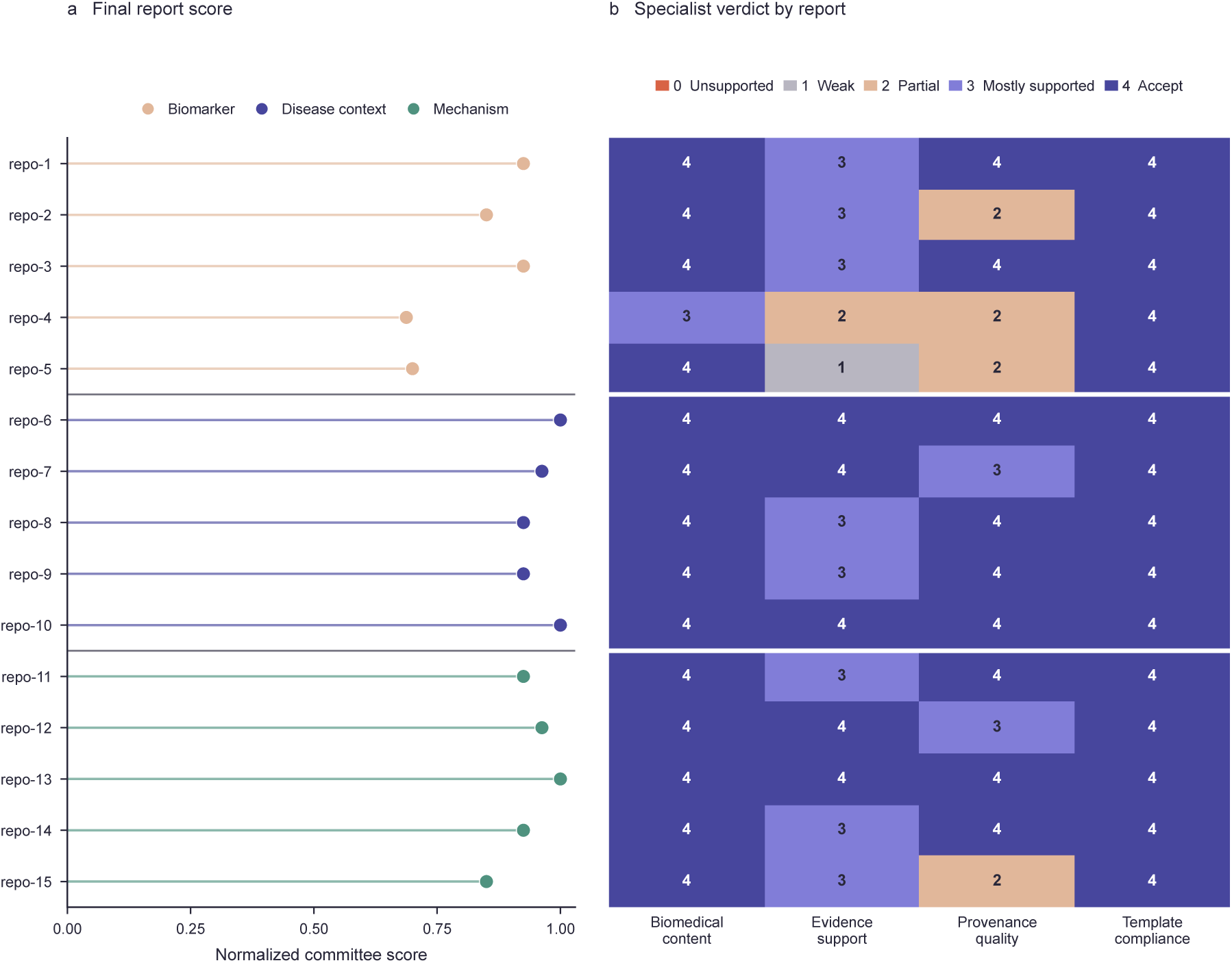
GeneReportBench report scores and specialist ratings. **a**, Normalized committee scores for all 15 reports, grouped by report type and shown on the complete zero-to-one scale. Marker colour distinguishes report types. **b,** Aligned specialist ratings for biomedical content, evidence support, provenance quality and template compliance. Levels range from 0 for Unsupported to 4 for Accept. Horizontal separators divide the three report types. All 15 runs completed and produced a saved report with a valid committee score.

### GLESKit produced answers for all human BixBench questions, with positive credit in most scored cases

Across the 67 selected BixBench questions, the system produced a final answer for 58. Of these answers, 56 received valid committee scores and two encountered judge failures, while nine questions had no answer. Nine valid scores were zero and 47 were positive. The valid score mean was 0.374 across 56 questions with a 95% CI from 0.287 to 0.461. The conservative mean was 0.322 across 65 questions. The positive only mean was 0.446 across 47 questions with a 95% CI from 0.355 to 0.536. Figure 4a presents the outcome counts. Figure 4b presents the three score summaries, with scope accounting in Extended Data Table 3.

The *H. sapiens* subset best matches the present operating domain. Of 41 questions, 39 received valid committee scores. Of these, 35 were positive and four were zero. The valid score mean and conservative mean were both 0.438, and the positive only mean was 0.488 (Fig. 4b). Across all 58 answered questions, including the two judge failures, the evidence specialist assigned the highest mean normalized rating at 0.431, followed by biology at 0.358 and statistics at 0.328 (Fig. 4c). The system therefore received at least partial credit for most validly scored *H. sapiens* questions, with statistical and numerical adequacy emerging as the weakest evaluated dimension. Review of zero score outcomes additionally identified numerical mismatch, trace inconsistency, biological misinterpretation, unsupported claims and incomplete execution (Extended Data Fig. 5).

**Fig. 4.**
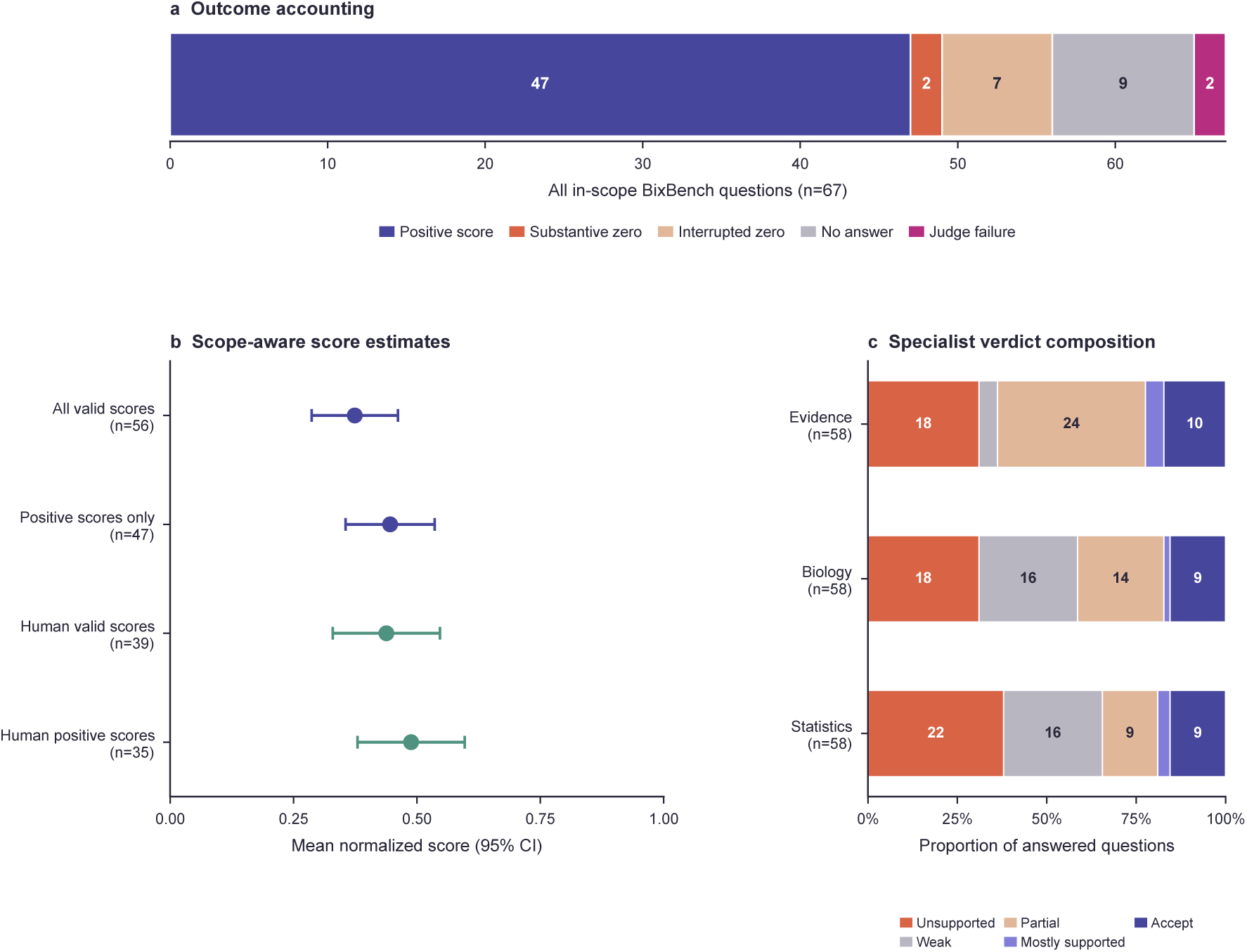
BixBench outcome accounting and performance. **a**, Outcome counts across all 67 selected questions. **b,** Points show valid score and positive only means and error bars show two sided 95% confidence intervals calculated with the Student’s *t* distribution and the sample standard error. Labels give the number of questions included in each mean. The plot label “Human” denotes the *H. sapiens* question set. **c,** Ordered rating categories from the statistics, biology and evidence specialists across all 58 answered questions. Numbers are shown for segments containing at least five verdicts.

### The human BixBench set showed higher completion and scores than the unmatched nonhuman sets

The BixBench question sets differed by species and case composition. The *H. sapiens* set contains 41 questions from 13 cases, the *M. musculus* set 13 questions from three different cases and the *P. aeruginosa* set 13 questions from four different cases. The valid score means were 0.438 across 39 *H. sapiens* questions, 0.206 across eight *M. musculus* questions and 0.246 across nine *P. aeruginosa* questions (Fig. 5a). The conservative means were 0.438, 0.127 and 0.170. All nine no answer outcomes occurred in the latter two sets, with five among the *M. musculus* questions and four among the *P. aeruginosa* questions (Fig. 5b). The observed differences describe question sets that vary in both species and case composition.

**Fig. 5.**
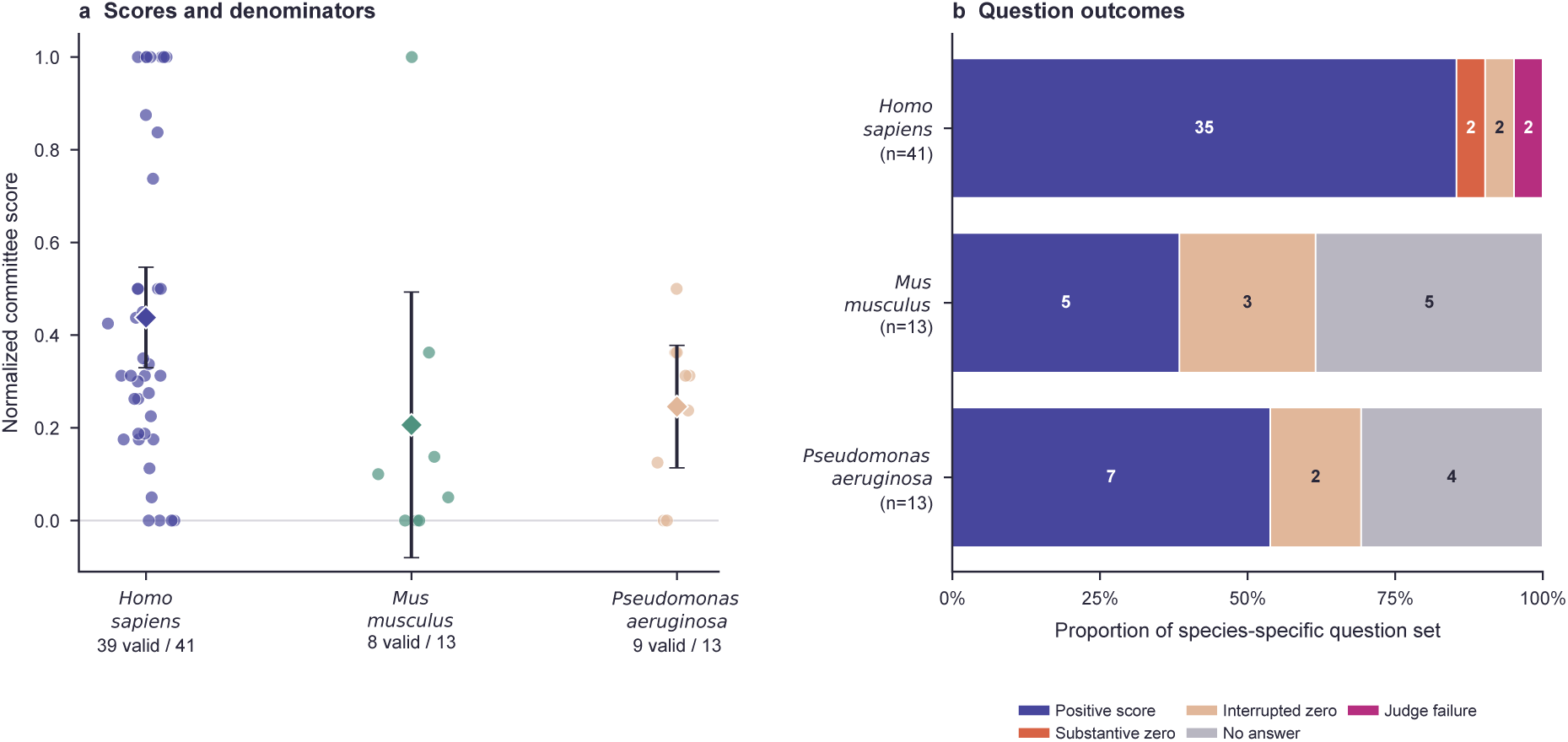
Performance across species specific BixBench question sets. **a**, Individual normalized committee scores for each question set. Diamonds show valid score means and error bars show two sided 95% confidence intervals calculated with the Student’s *t* distribution and the sample standard error. Species labels give the number of valid scores and total questions. **b,** Outcome counts within each full species set. Species and case composition are confounded because the three sets contain different cases.

### Configurations with skills scored higher in both paired evaluations and made more tool calls

In the primary paired analysis of the same 20 *H. sapiens* BixBench questions, the mean committee score was 0.494 with skills and 0.349 without skills, giving a mean paired difference of 0.145 (*P* = 0.103, Fig. 6b). Thirteen questions improved, four were unchanged and three worsened. The configuration with skills completed 19 episodes compared with 17 without skills and made 196 total tool calls compared with 89 (Fig. 6a). In the valid pair sensitivity analysis of 19 questions, mean scores were 0.517 with skills and 0.367 without skills, with a paired difference of 0.150 (*P* = 0.111). Twelve questions improved, four were unchanged and three worsened. In the completed valid sensitivity analysis of 16 pairs, the means were 0.489 and 0.436, with a difference of 0.053. The positive score difference persisted in both sensitivity analyses. Completion and answer quality remain combined in these selected subsets. The paired scores and completion statuses are supplied in the Source Data file.

In the corresponding paired comparison of five GeneReportBench reports, the mean committee score was 0.925 with skills and 0.840 without skills, giving a mean within report difference of 0.085 (*P* = 0.125, Fig. 6d). Four reports improved, one was unchanged and none worsened. The configuration with skills made 130 total tool calls compared with 95 without skills (Fig. 6c). Across these five reports, the direction of the score difference matched the BixBench comparison. Both comparisons are descriptive of these task sets and configurations. These observations suggest that skills may support more effective use of the available tools through task specific guidance.

**Fig. 6.**
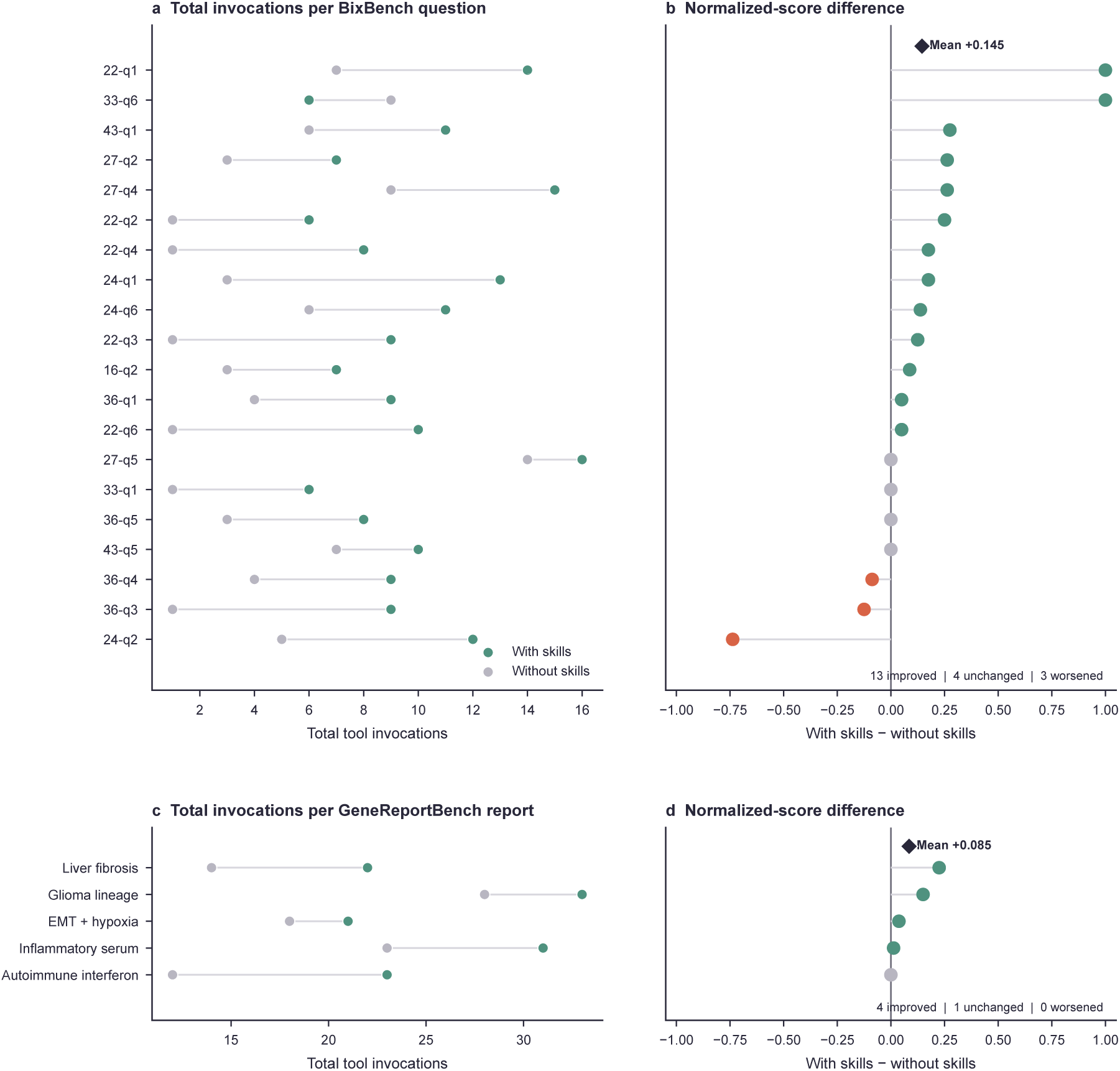
Paired skills comparisons for focused questions and extended reports. **a**, Total tool calls for 20 BixBench questions with and without skills. **b,** Paired normalized score difference for the same questions, calculated as performance with skills minus performance without skills. **c,** Total tool calls for five GeneReportBench reports with and without skills. **d,** Paired normalized score difference for the same reports. Each tool call is counted separately, including repeated calls to the same tool. Connected circles identify paired configurations. In **b** and **d**, green circles indicate improvement, coral circles indicate worsening and grey circles indicate no change. Diamonds denote mean paired differences.

## Discussion

Recent systems establish complementary precedents for scientific agents. Biomni [12] demonstrates broad biomedical task composition within a unified action environment, while the agentic X-ray scientist [13] shows that existing reasoning models coupled to MCP tools and persistent state can conduct a closed-loop physical workflow. GLESKit develops a distinct configuration for HITL gene-level investigation. It separates a modular tool layer from the model and conversational client, makes the same operations available through compatible access routes, and connects an optional persistent workspace to evidence-linked analysis and initial report production. This configuration allows biomedical researchers to direct an investigation through a conversational environment while retaining outputs for inspection and revision. The contribution is an integrated, gene-centred workflow that combines portable biomedical operations, persistent study materials and structured evidence synthesis.

The two benchmarks provide complementary evidence about this environment. GeneReportBench shows that the evaluated workflow can complete controlled, extended tasks and retain structured reports with linked materials. Its specialist assessments separate structural completion from evidence quality by revealing weaker evidence and provenance in some reports despite strong template compliance and biomedical content. BixBench provides a stricter view of focused answer quality and identifies statistical and numerical reliability as immediate constraints. Relevant evidence supports source tracing and failure diagnosis. An incorrect quantitative conclusion still constitutes a failure. Together, the results show that GLESKit can organize a substantive investigation, earn positive credit on most validly scored gene-level questions about human biology and retain materials for expert assessment. Its strongest demonstrated use is therefore the organization of gene-level investigations of human biology whose evidence and calculations remain available for review.

The paired evaluations provide the clearest comparison of the configurations. Comparing matched focused questions and extended reports with and without skills showed differences in scores and tool use while matching the model, tool access, MCP service and scoring procedure. Scores and tool use were higher with skills in both task types, although the GeneReportBench comparison included only five reports and the BixBench comparison used different OpenCode versions between conditions. The BixBench score difference remained when the analysis was restricted to questions with valid scores in both configurations. These differences apply to the evaluated task sets and configuration.

The evidence characterizes the configured GLESKit system rather than establishing superiority over systems such as GeneGPT or Biomni, whose tasks, interfaces and available resources differ. GeneReportBench contains 15 author-curated tasks, while the BixBench analysis uses a scope-filtered subset. Both evaluations rely on model-based specialist judges and contain no human ratings. The analysis used one retained outcome per task and configuration rather than repeated independent runs. The confidence intervals therefore describe variation across tasks rather than variation across repeated executions. The HITL workflow is an intended mode of use, not an outcome tested by these automated benchmarks. Session-linked artifacts support inspection, while the sufficiency and interpretation of their evidence remain responsibilities for expert review.

The species-specific BixBench results also require proper interpretation. The human, mouse and bacterial questions came from different cases, so species is confounded with task composition and cannot explain the score differences on its own. The concentration of no-answer outcomes in the mouse and bacterial sets describes the current evaluated scope. It does not estimate an isolated species effect. A shared gene identifier can organize further modalities and organisms, but each addition requires suitable resources, mappings, methods and validation.

## Methods

### Operational terminology

We distinguish the generative model from the system that surrounds it. A large language model (LLM) is the model producing and interpreting language. An agent uses an LLM to maintain task context and select tools in response to a request. The agent runtime is the execution environment that combines model access, permissions, skills, tools and control flow. In the GLESKit web application, OpenCode [21] supplies this general-purpose runtime. A skill provides reusable instructions, constraints or decision heuristics for a class of task. A tool is a callable retrieval, analysis, visualization or reporting function. A biomedical operation is a user-facing capability implemented by one or more tools. A tool call (labelled an invocation in the figures) is one execution of a tool, whereas a distinct-tool count records the number of unique tools used. MCP [19] is the client–server protocol through which tools are described, discovered and invoked. These definitions separate model behaviour from capabilities supplied by external computation and orchestration.

A study session is the persistent unit used by the web application to organize one investigation. An artifact is a stored intermediate or final research output associated with a session, including uploaded data, a note, structured analysis, evidence bundle, figure or report. In this operational context, artifact denotes a stored research object. Biological artifacts refer separately to noise, bias or technical defects. A provider is a local analytical component or external data service accessed behind a GLESKit tool. A report template is a modifiable Markdown specification that defines the expected sections and content guidance for a class of report while leaving the scientific conclusions to the investigation.

### System scope and gene-centred organization

GLESKit accepts several research entry points, including a gene or a gene set, raw RNA-sequencing counts, differential-expression results, a biomedical question or a hypothesis. Its current operations cover identifier resolution, gene and protein characterization, expression profiling, differential-expression and gene-set enrichment analyses, and pathway or interaction-network interpretation. Further operations retrieve gene–disease, variant–trait and drug–target context, gather literature, generate figures and support report production (Extended Data Table 5). Providers return typed results that retain the source and analytical context. Shared components normalize gene identifiers and convert provider outputs into forms that can be passed to another tool or stored in the workspace.

A complete inventory of the tools available in GLESKit, including their purposes and supporting providers or local engines, is provided in Supplementary Table 1.

The system uses genes as the common entity for relating results from transcriptomics, protein annotation, human genetics and pharmacogenomics. Provider-specific results preserve their modality and evidentiary type, while normalized gene identifiers allow them to be assembled around the same investigation. A new resource or analytical operation requires an appropriate provider adapter and validation, while the client-facing MCP interface remains stable. A new modality additionally requires a biologically justified mapping to gene-level entities and modality-specific validation. These extension points implement modularity while assigning each new tool or modality its own validation requirements.

### MCP architecture and tool execution

The implementation separates a client layer, an MCP access layer and a backend provider layer (Extended Data Fig. 1). The local MCP service uses STDIO. The MCP service can optionally be exposed to compatible remote hosts using the implementation’s SSE transport [22]. This optional network connection is separate from the locally served workspace interface. The backend server declares the available tools and delegates calls through an orchestrator to external services, local analytical functions and shared normalization or formatting components. Local tools support workspace management, data preparation, statistical analysis and visualization; examples include session storage, Principal Component Analysis (PCA) and DESeq2 [23]. External providers supply annotation, pathway, interaction, association and literature records. Integrated workflows can combine both: identifier resolution uses local references with remote fallbacks, and enrichment can combine local R computation with external pathway information (Extended Data Table 5). Execution location and scientific function therefore describe different aspects of a tool. Provider adapters can be added behind this layer while preserving how compatible clients discover the operations. Provider-specific records retain their type and source when identifiers and outputs are normalized. The benchmarks used the configured MCP route.

The web chat interface uses an OpenCode server [21] to instantiate the agent, connect configured model providers, load skills and manage tool calls. This separates model selection from the biomedical MCP service.

### Workspace and report lifecycle

Users launch the web application on their own computer and open its local address in a browser to access the workspace and chat. The workspace interface is served locally, not hosted as a public website. Sessions, uploaded files and generated artifacts are stored on the user’s computer, separately from the transient model context. Tool results and user-supplied materials can be saved as artifacts with identifiers linking them to their session and type. Subsequent operations can retrieve these objects for revision, visualization, comparison or synthesis. Report generation uses modifiable Markdown templates. The three templates evaluated here were Biomarker Prioritization, Mechanism Interpretation and Disease Context. Each defines required sections and gives task-specific guidance while reserving scientific conclusions for the investigation. The resulting report is itself saved as an artifact and can link the intermediate materials used during its preparation.

The intended workflow keeps the HITL throughout this lifecycle. The agent proposes or executes bounded operations, while the user may inspect outputs, change the direction of the analysis and decide what is included in the report. Reports serve as initial, inspectable syntheses for expert interpretation. The controlled benchmark runs were automated and evaluated system-generated outputs without HITL interventions.

### GeneReportBench design and scoring

GeneReportBench, developed for this study, is an author-curated benchmark of structured biomedical report production. It contains 15 tasks about human biology divided equally among biomarker prioritization, mechanism interpretation and disease context. Each instance comprises a controlled prompt, a corresponding report template, four expected content nuggets and a list of prohibited overclaims. Expected nuggets are task-specific factual or analytical elements that a complete report should address. Prohibited overclaims are statements that would go beyond the evidence allowed by the task. The expected nuggets and prohibited overclaims guided the specialist assessments. The committee score was calculated from the four specialist ratings described below. The evaluated tasks and report type definitions are provided in the evaluation archive described in Data availability. A benchmark-specific agent received bounded access to the relevant GLESKit skills and MCP tools and was required to save the final report in the task’s study session.

Four specialist roles assessed biomedical content, evidence support, provenance quality and template compliance in the saved report. Each specialist assigned a rubric-anchored rationale and an ordered level. Unsupported (0) denotes no adequate support or a materially incorrect result. Weak (1) denotes limited support with major deficiencies. Partial (2) denotes some relevant support but material omissions or weaknesses. Mostly Supported (3) denotes substantial support with minor limitations. Accept (4) denotes that the criterion was met without a material deficiency. The four levels were normalized to the 0–1 interval and combined with weights of 0.350 for biomedical content, 0.300 for evidence support, 0.150 for provenance and 0.200 for template compliance. These weights sum to one, so the committee score is their direct weighted mean with no additional cap or penalty. An arbiter synthesized the four specialist assessments qualitatively; it did not supply a fifth weighted rating. All specialist and arbiter calls used MiniMax M3 through OpenRouter at temperature zero, with the agent model selected through the procedure described below. The judge context included the saved report, task rubric, report metadata, nugget overlap and provenance values and compacted transcript and tool-provenance summaries. Evidence and provenance specialists also received an offline citation audit. They therefore assessed supplied session evidence as well as the report text; a count of lines matching a prescribed Sources format was not a count of all available citations. High aggregate scores can coexist with weaker evidence or provenance because the weighted mean also includes content and template compliance.

The example in Fig. 2b also displays nugget overlap and provenance values. Nugget overlap is the mean RapidFuzz partial string-match similarity between each expected nugget and the report, normalized to 0–1. It measures textual overlap only. The provenance value combines counts of recognized source lines, provider-attributed lines, Markdown links and embedded figures, with weights 0.25, 0.35, 0.25 and 0.15 and saturation counts 4, 3, 4 and 1, respectively. It subtracts 0.05 per source line containing an opaque artifact identifier, up to 0.25, and is bounded below by zero. These values were supplied to all four specialists. The committee score was calculated from the four categorical ratings, with no separate weighting for these values.

For each episode, we recorded completion, whether the final report was saved, report length, total tool calls and linked artifacts. A total tool-call count includes repeated calls to the same tool, whereas a distinct-tool count includes each tool only once. These values describe workflow activity and do not contribute to the committee score. Quality outcomes and retained workflow measures for each task are shown in Extended Data Tables 1 and 2. The same records are supplied in machine readable form in the Source Data file.

### BixBench scope, execution and scoring

BixBench [20] contains open-answer computational-biology tasks spanning multiple data modalities and analytical requirements. Before full evaluation, we defined an allowlist for questions involving gene-expression matrices, bulk or single-cell RNA analysis, differential expression, enrichment, clustering, supervised modelling or gene-centred reporting. We excluded questions whose main requirement concerned variants, whole-genome sequencing, imaging, proteomics-first or epigenomics-first analysis, or clinical modelling without an RNA-centred component. Eighteen complete cases and two question-level exceptions from mixed cases met these criteria, reducing 205 questions to 67 from 20 benchmark cases. The selected questions comprised 41 concerning *H. sapiens* from 13 cases, 13 concerning *M. musculus* from three cases and 13 concerning *P. aeruginosa* from four cases. The species sets contain different cases and questions.

Each question was supplied to a benchmark-specific agent with a task-specific role prompt and bounded GLESKit skills and MCP tools. Statistics, biology and evidence specialists then assessed the returned answer against the benchmark reference, followed by a qualitative arbiter. Numeric questions used weights of 0.550, 0.200 and 0.250 for statistics, biology and evidence, respectively. Semantic questions used weights of 0.250, 0.400 and 0.350. In each scheme the weights sum to one. The ordered specialist levels were normalized to 0–1, and the committee score was their direct weighted mean with no additional post hoc cap or penalty. For the manuscript analysis, these weighted means were recomputed from persisted specialist categories without rerunning judges; they can differ from originally stored aggregate scores subject to earlier aggregation policies. A score of 1.000 was therefore the maximum. The arbiter supplied a qualitative interpretation of the specialist result.

Species specific summaries describe the observed question sets because species and case composition are confounded. Scope accounting is shown in Extended Data Table 3 and supplied in machine readable form in the Source Data file.

### BixBench outcome classes and score denominators

A positive score is a valid committee score greater than zero. A substantive zero is a valid zero score attached to a completed final answer. An interrupted zero is a valid zero score attached to retained partial output or an interrupted execution. A no answer outcome has no final answer and no committee score. A judge failure has an answer but no valid committee verdict. Both zero categories are valid scores and remain included whenever an analysis uses valid committee scores. For the full cohort, the valid score mean includes valid zeros and excludes no answer outcomes and judge failures. The conservative cohort mean includes valid zeros, assigns zero to no answer outcomes and excludes judge failures. The positive only mean includes exclusively valid scores greater than zero.

The primary BixBench skills comparison retains all 20 assigned questions per configuration. An episode without a valid verdict, including a judge failure, is assigned zero because completion is part of the comparison. The skills sensitivity analyses use 19 pairs with valid scores in both configurations and 16 pairs that both completed and received valid scores. All GeneReportBench observations were scored and nonzero, so these denominator choices do not change its reported means. Outcome treatment and denominators across the reported analyses are summarized in Extended Data Table 4.

### Model selection and paired skills ablations

We used two budget-constrained paired comparisons to select the agent model. Terra was accessed through OpenAI and DeepSeek through OpenRouter. On 20 BixBench questions, GPT-5.6 Terra achieved a mean normalized score of 0.593 and DeepSeek V4 Flash 0.494. Terra scored higher on ten questions, DeepSeek on three and seven were equal. Terra’s three largest gains accounted for 73.7% of the total positive score difference (Extended Data Fig. 2). All 20 questions concerned *Homo sapiens*. Their source-provided BixBench categories were multi-label and overlapping, with Transcriptomics assigned to 18 questions, RNA-seq to 17, Sequence Analysis to 12, Differential Expression Analysis to 11, Machine Learning and Artificial Intelligence to three, Single-Cell Analysis to two and Functional Genomics to one. Category-level score differences are descriptive because the labels overlap and several groups are small (Extended Data Fig. 3). Based on token counts and OpenRouter prices recorded at evaluation time, Terra’s aggregate inference cost was approximately eight times higher. On five paired GeneReportBench tasks, the corresponding mean scores were 0.875 for Terra and 0.925 for DeepSeek V4 Flash, with DeepSeek higher on four reports, while Terra cost approximately 8.7 times more (Extended Data Fig. 4). These samples supported only the resource allocation decision. DeepSeek V4 Flash was retained for the complete evaluations. The evaluation sequence and reuse of benchmark runs are summarized in Extended Data Fig. 6.

After selecting DeepSeek V4 Flash, we evaluated skills on the same 20 BixBench questions and five GeneReportBench tasks used for model selection. The original DeepSeek runs with skills served as baselines for subsequent runs without skills. The BixBench mean of 0.494 consequently describes the same run in both comparisons. Pairing denotes comparison within each task under the two configurations. Within each paired comparison, the agent model, MCP service, exposed tool schema and scoring procedure were matched. The condition without skills used corresponding role and task prompts that removed skill loading and skill specific guidance while retaining the same benchmark item and tool access. We recorded the final committee score and total tool calls for both comparisons and BixBench completion status. Each call is counted separately, including repeated calls to the same tool. The reported paired difference is the score with skills minus the score without skills and applies only to each evaluated task set and configuration.

### Execution settings

BixBench runs were limited to 12 assistant turns, 30 tool calls and 1,200 seconds per question, whereas GeneReportBench runs were limited to 14 turns, 36 tool calls and 900 seconds per task. Each task and configuration contributed one retained outcome, including partial output from interrupted runs when it could be judged. Reported tool counts include skill loading, file reading and biomedical MCP calls. The BixBench skills conditions used different OpenCode releases, making the runtime version a potential contributor to the observed difference. Complete execution settings are provided in the evaluation archive described in Data availability.

### Statistical summaries

Scores are summarized on their normalized 0–1 scale. We report arithmetic means, and for the main GeneReportBench summary additionally the median, sample standard deviation and range. Two sided 95% confidence intervals around means use the Student’s *t* distribution and the sample standard error. These intervals describe variation across the observed tasks. Reported *P* values use two sided exact paired permutation tests of the mean score difference between configurations.

### Generative artificial intelligence in manuscript preparation

OpenAI Codex and Prism were used to suggest and refine wording, organization and formatting during manuscript preparation. The authors reviewed and verified all changes and take responsibility for the final text.

## Data availability

The study data comprise benchmark evaluation records. Source data supporting the quantitative figures and tables are provided with this paper. Final evaluated outputs, final judging records, task definitions, retained execution records and evaluation configuration are available in the evaluation archive at https://doi.org/10.5281/zenodo.23592806 (ref. [24]). The BixBench benchmark is described in ref. [20].

## Code availability

The GLESKit source code, technical documentation and analysis notebook used to generate the manuscript figures and tables are available at https://github.com/4th05/GLESKit. The documentation includes the mapping between tools and providers.

## Acknowledgements

A.M. acknowledges support from the Fundação para a Ciência e a Tecnologia, I.P. (FCT), through a doctoral scholarship, and from INESC TEC as the host institution.

## Author contributions

Athos Moraes conceived the study, designed and implemented GLESKit, designed and conducted the evaluations, analysed the results and wrote the original manuscript. Pedro Ferreira supervised the work, co-conceived the study, contributed to its conceptual development, interpreted the results and revised the manuscript. Both authors discussed the findings and approved the final manuscript.

## Extended Data

**Extended Data Fig. 1.**
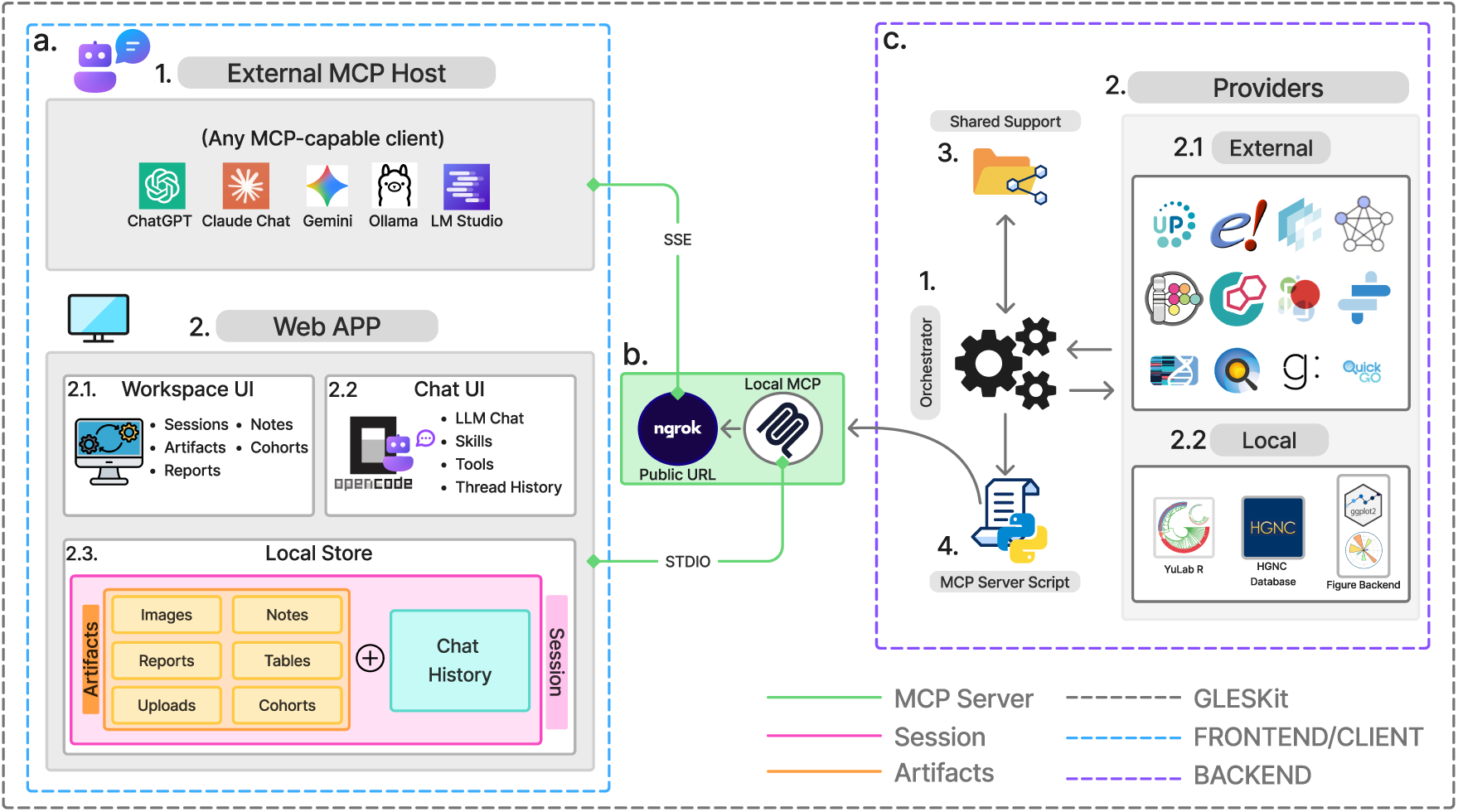
Internal GLESKit architecture. **a,** External MCP hosts and the locally run web application’s workspace, chat and local-store components, with browser access to the workspace and chat. **b,** MCP access over local STDIO or optional networked SSE; the network connection exposes the MCP service, not the workspace web interface. **c,** Backend orchestration across external providers, local analytical providers, shared components and the MCP server declaration.

**Extended Data Fig. 2.**
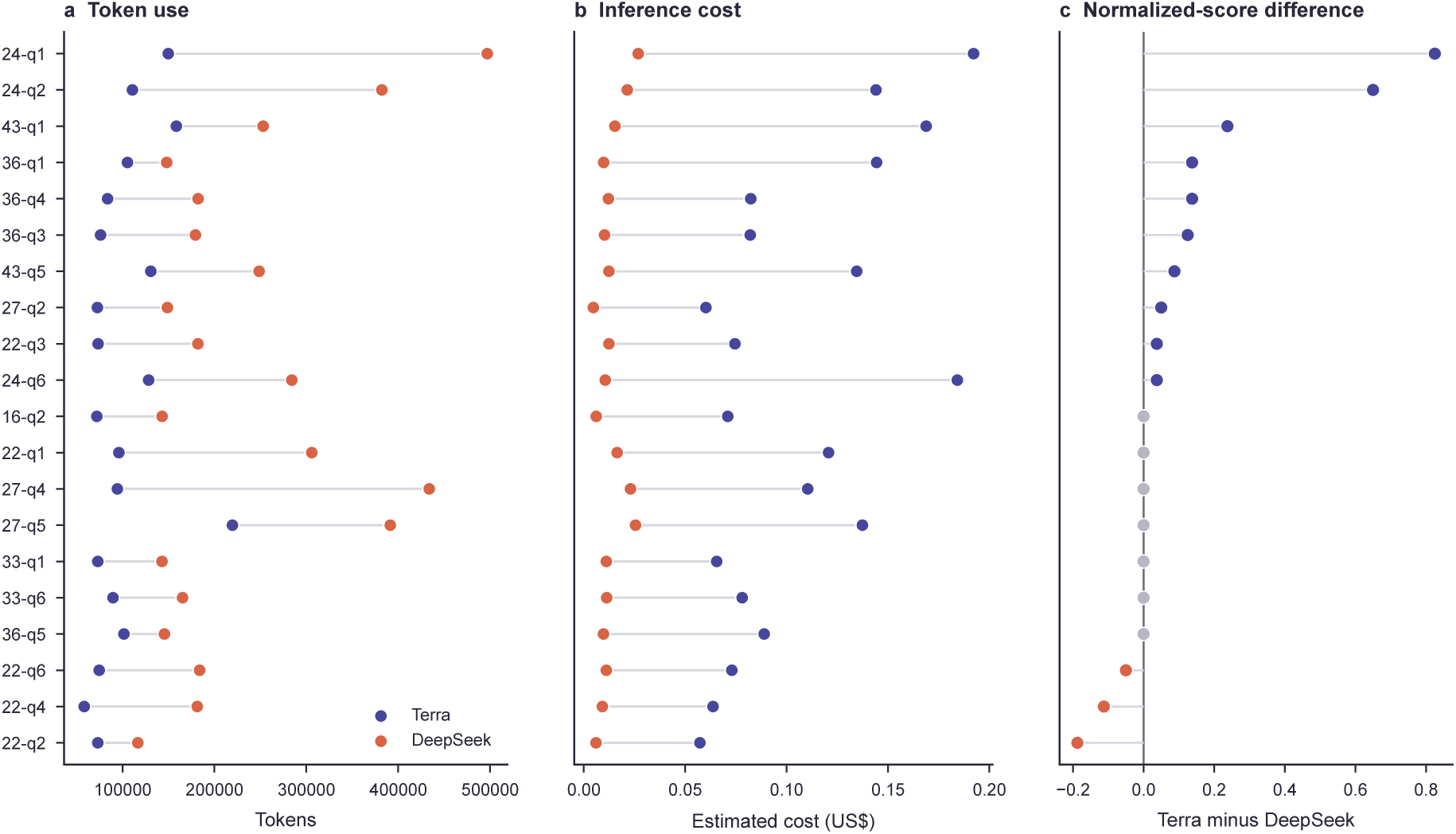
Paired BixBench resource use, cost and score differences. **a**, Reported token use. **b,** Estimated inference cost. Connected markers in **a** and **b** identify the paired Terra and DeepSeek observations for each of 20 questions. **c,** Paired normalized score difference calculated as Terra minus DeepSeek. Positive differences favour Terra.

**Extended Data Fig. 3.**
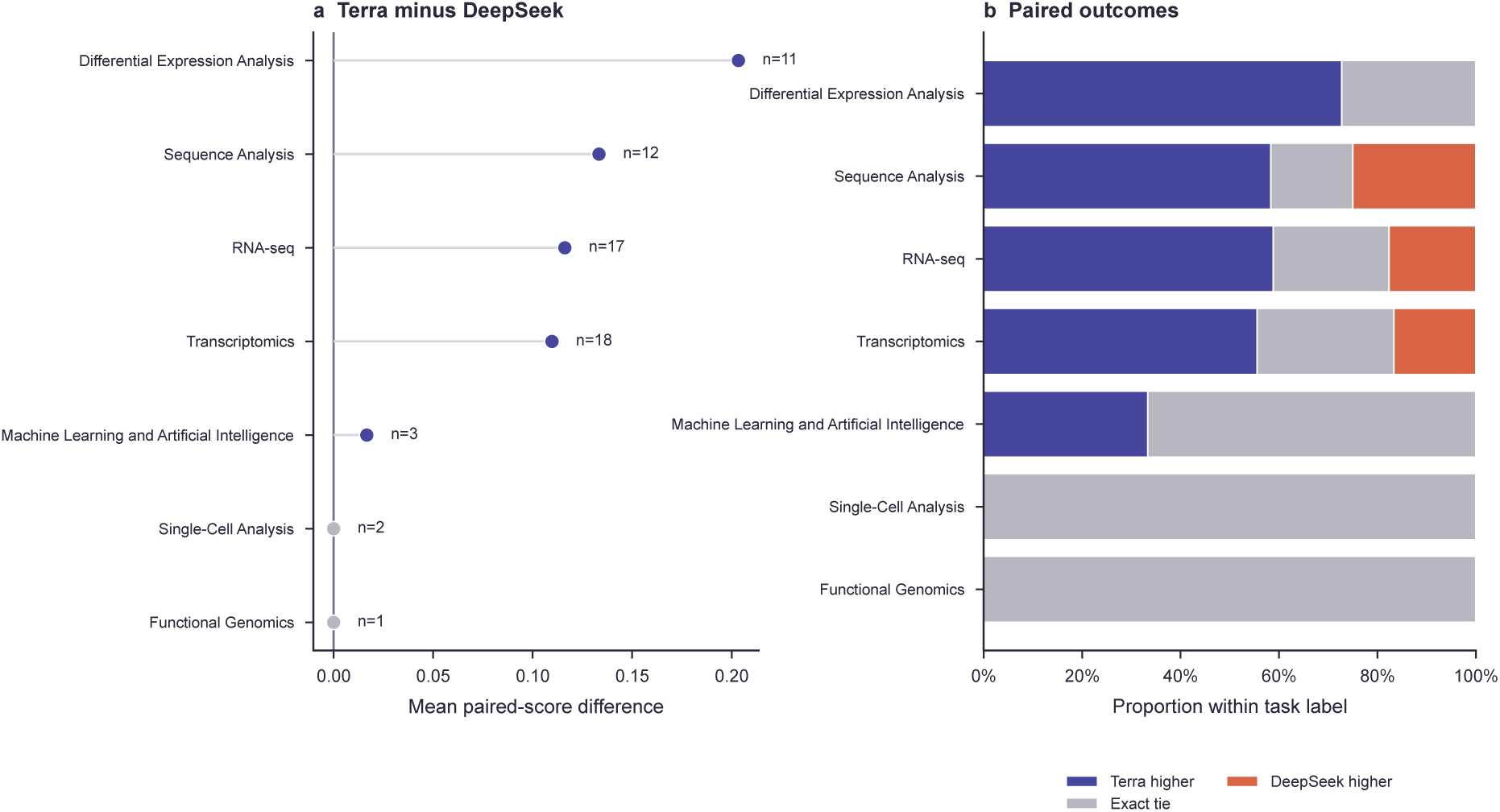
Metadata-driven BixBench task-category analysis of the paired model-selection questions. **a**, Mean paired normalized-score difference between GPT-5.6 Terra and DeepSeek V4 Flash within each BixBench task category. Points and stems show the category estimates relative to zero. Positive values favour Terra and negative values favour DeepSeek. Labels give the number of questions carrying each category. **b,** Proportions of Terra wins, exact ties and DeepSeek wins within each category. All 20 questions concern *Homo sapiens*. Categories are source-provided multi-label annotations, so their counts overlap and do not sum to 20.

**Extended Data Fig. 4.**
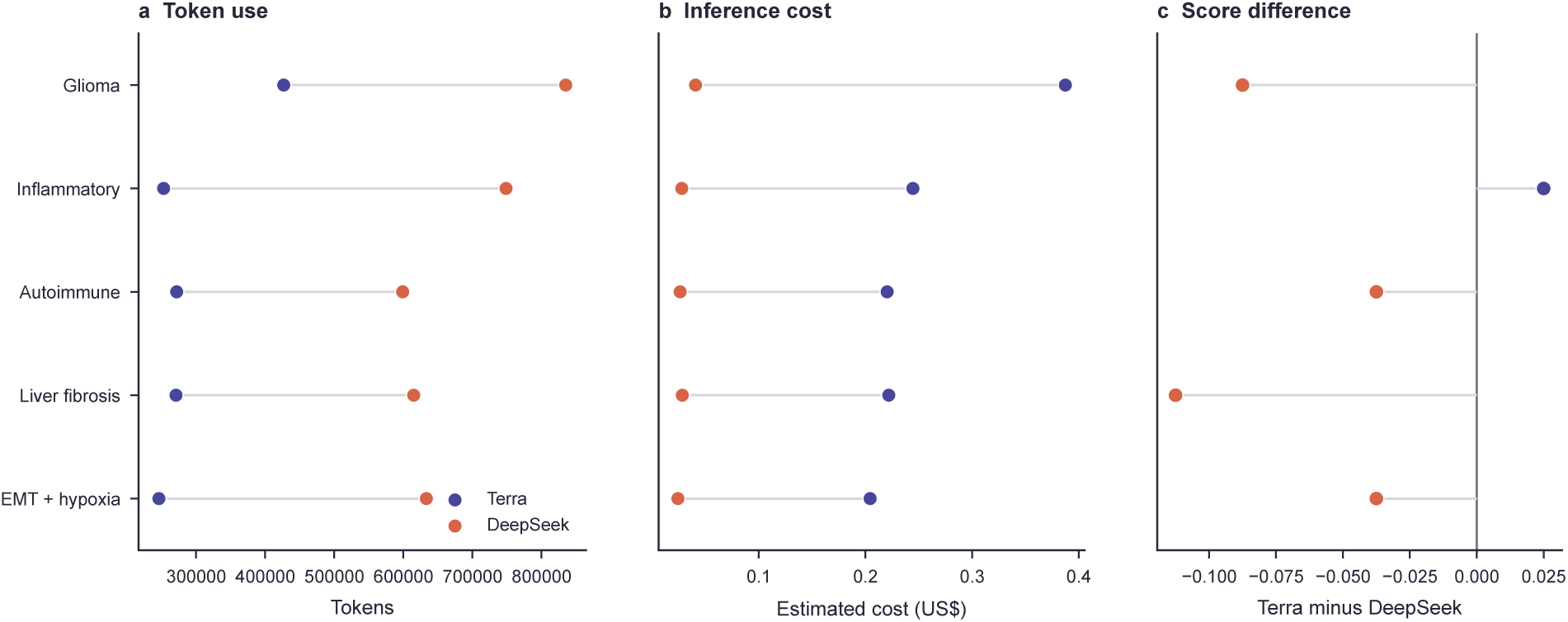
Paired GeneReportBench model-selection results. **a**, Reported token use. **b,** Estimated inference cost. Connected markers in **a** and **b** identify paired Terra and DeepSeek observations across five reports. **c,** Paired committee-score difference calculated as Terra minus DeepSeek. Positive differences favour Terra and negative differences favour DeepSeek.

**Extended Data Fig. 5.**
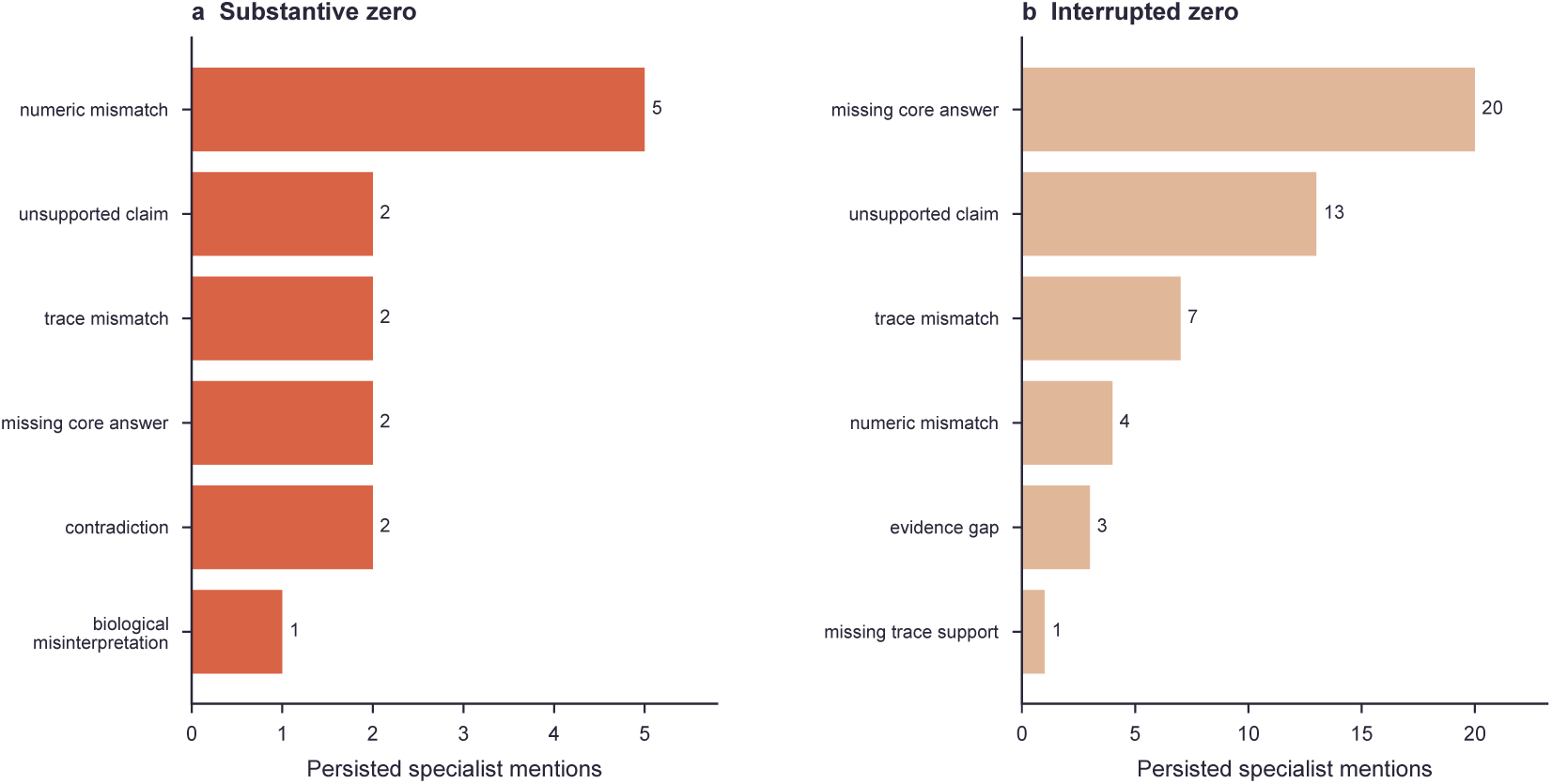
Specialist findings associated with BixBench zero score outcomes. **a**, Findings associated with substantive zero score answers. **b,** Findings associated with interrupted zero score answers. The panels summarize two substantive zero outcomes and seven interrupted zero outcomes. Bar end labels give persisted specialist mentions, and more than one finding may be recorded for a question.

**Extended Data Fig. 6.**
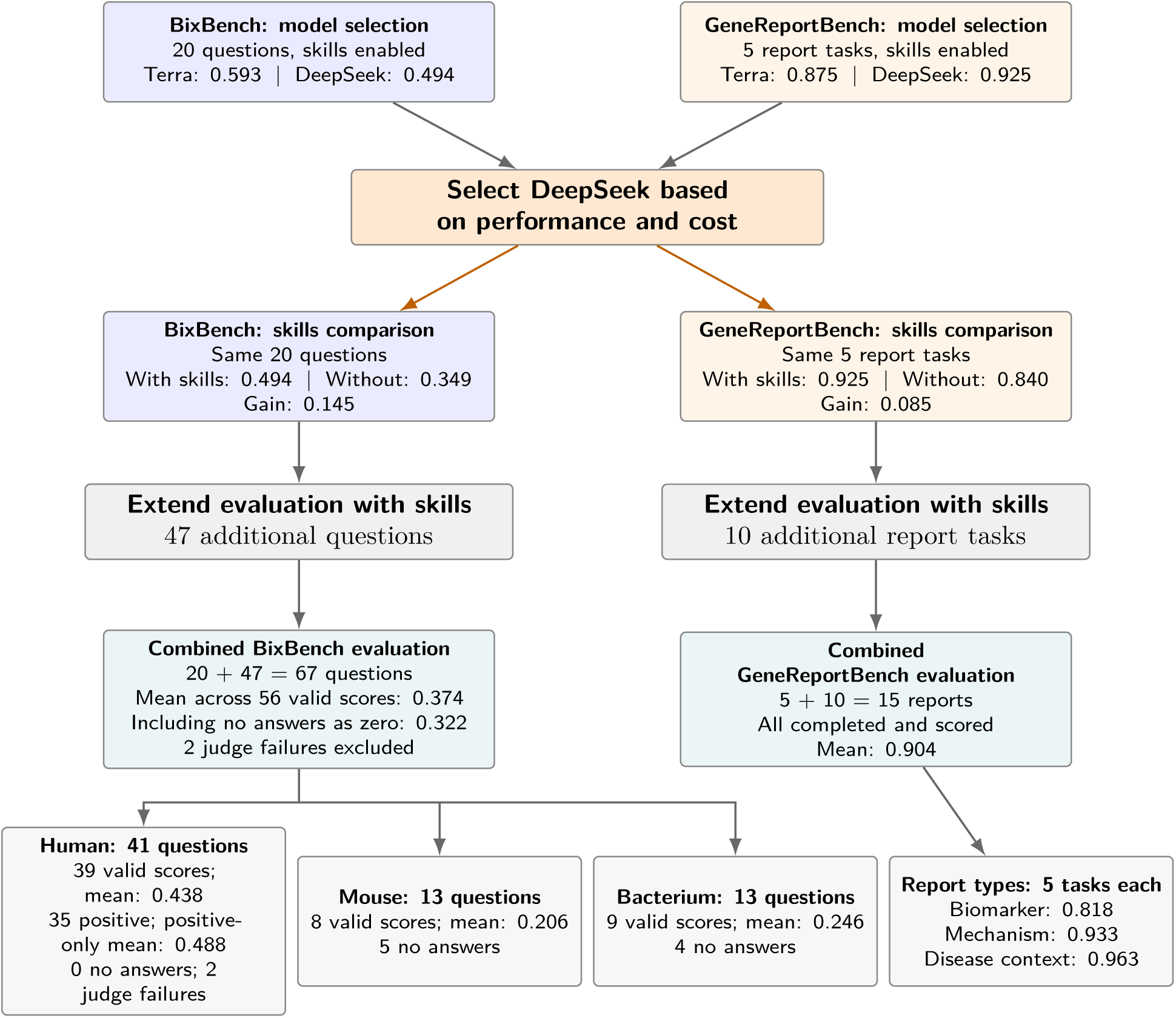
Evaluation sequence and reuse of benchmark runs. DeepSeek and Terra were first compared on 20 BixBench questions and five GeneReportBench tasks. After selecting DeepSeek, we compared performance with and without skills, then expanded the evaluation to 67 questions and 15 reports. The diagram gives the task counts and mean scores for each comparison and the final benchmark summaries.

**Extended Data Table 1.** GeneReportBench task-level quality outcomes. All reports were completed, saved and scored. Specialist levels use 0 for Unsupported, 1 for Weak, 2 for Partial, 3 for Mostly Supported and 4 for Accept. B denotes biomedical content, E evidence support, P provenance quality and T template compliance.

| Task | Report type | Final | B | E | P | T |
| --- | --- | --- | --- | --- | --- | --- |
| repo-1-q1 | Biomarker | 0.925 | 4 | 3 | 4 | 4 |
| repo-2-q1 | Biomarker | 0.850 | 4 | 3 | 2 | 4 |
| repo-3-q1 | Biomarker | 0.925 | 4 | 3 | 4 | 4 |
| repo-4-q1 | Biomarker | 0.688 | 3 | 2 | 2 | 4 |
| repo-5-q1 | Biomarker | 0.700 | 4 | 1 | 2 | 4 |
| repo-6-q1 | Disease context | 1.000 | 4 | 4 | 4 | 4 |
| repo-7-q1 | Disease context | 0.963 | 4 | 4 | 3 | 4 |
| repo-8-q1 | Disease context | 0.925 | 4 | 3 | 4 | 4 |
| repo-9-q1 | Disease context | 0.925 | 4 | 3 | 4 | 4 |
| repo-10-q1 | Disease context | 1.000 | 4 | 4 | 4 | 4 |
| repo-11-q1 | Mechanism | 0.925 | 4 | 3 | 4 | 4 |
| repo-12-q1 | Mechanism | 0.963 | 4 | 4 | 3 | 4 |
| repo-13-q1 | Mechanism | 1.000 | 4 | 4 | 4 | 4 |
| repo-14-q1 | Mechanism | 0.925 | 4 | 3 | 4 | 4 |
| repo-15-q1 | Mechanism | 0.850 | 4 | 3 | 2 | 4 |

**Extended Data Table 2.** GeneReportBench retained workflow materials. Linked artifacts are session objects associated with the saved report. Total tool-call counts include repeated calls to the same tool. These workflow measures do not contribute to the committee score.

| Task | Short title | Artifacts | Words | Total tool calls |
| --- | --- | --- | --- | --- |
| repo-1-q1 | Epithelial tumor biomarker panel | 11 | 3738 | 31 |
| repo-2-q1 | Inflammatory serum marker short-list | 16 | 2761 | 31 |
| repo-3-q1 | Glioma lineage and invasion panel | 18 | 3686 | 33 |
| repo-4-q1 | Fibrosis extracellular matrix panel | 11 | 2680 | 28 |
| repo-5-q1 | Cardio-inflammatory biomarker shortlist | 13 | 3151 | 26 |
| repo-6-q1 | Inflammatory bowel disease panel | 12 | 2623 | 22 |
| repo-7-q1 | Microglial neurodegeneration panel | 7 | 2306 | 21 |
| repo-8-q1 | Pulmonary fibrosis panel | 11 | 2538 | 25 |
| repo-9-q1 | Autoimmune interferon panel | 13 | 2270 | 23 |
| repo-10-q1 | Liver fibrosis and susceptibility panel | 9 | 1934 | 22 |
| repo-11-q1 | EMT and hypoxia program | 9 | 2162 | 21 |
| repo-12-q1 | Interferon response program | 9 | 2227 | 17 |
| repo-13-q1 | Cell-cycle and mitotic program | 10 | 2219 | 24 |
| repo-14-q1 | Stress and apoptosis program | 7 | 2534 | 17 |
| repo-15-q1 | Mitochondrial respiratory program | 12 | 2417 | 19 |

**Extended Data Table 3.** Scope aware BixBench outcome accounting. Counts and mean scores are shown for the full cohort and the three species sets. The species sets contain different cases and are not matched comparisons.

| Cohort | Total | Valid | Positive | Zero | No answer | Judge failure | Mean valid | Mean no-answer=0 |
| --- | --- | --- | --- | --- | --- | --- | --- | --- |
| All in-scope | 67 | 56 | 47 | 9 | 9 | 2 | 0.374 | 0.322 |
| *Homo sapiens* | 41 | 39 | 35 | 4 | 0 | 2 | 0.438 | 0.438 |
| *Mus musculus* | 13 | 8 | 5 | 3 | 5 | 0 | 0.206 | 0.127 |
| *Pseudomonas aeruginosa* | 13 | 9 | 7 | 2 | 4 | 0 | 0.246 | 0.170 |

**Extended Data Table 4.** Outcome treatment and denominators across reported analyses. The table shows how valid zero scores, no answer outcomes and judge failures are treated in each analysis. Mean denominators give the number of observations used for each reported mean.

| Analysis | Valid zero | No answer | Judge failure | Mean denominator |
| --- | --- | --- | --- | --- |
| BixBench model selection | Included | Not observed | Not observed | 20 per model |
| BixBench skills primary analysis | Included | Assigned zero if present | Assigned zero | 20 per configuration |
| BixBench skills valid pair sensitivity | Included | Excluded | Excluded | 19 pairs |
| BixBench skills completed valid sensitivity | Included | Excluded | Excluded | 16 pairs |
| Combined BixBench valid score mean | Included | Excluded | Excluded | 56 |
| Combined BixBench conservative mean | Included | Assigned zero | Excluded | 65 |
| Combined BixBench positive only mean | Excluded | Excluded | Excluded | 47 |
| GeneReportBench model selection | Included, none observed | Not observed | Not observed | 5 per model |
| GeneReportBench skills comparison | Included, none observed | Not observed | Not observed | 5 per configuration |
| Combined GeneReportBench evaluation | Included, none observed | Not observed | Not observed | 15 |

**Extended Data Table 5.** Preview of GLESKit tools and supporting resources. The table provides a preview of GLESKit tools, classified in four categories, and their supporting resources. The complete inventory of all 75 tools, including their purposes and provider or local engine mappings, is provided in Supplementary Table 1.

| Category | Function | Representative tools | Providers or engines |
| --- | --- | --- | --- |
| External evidence | Retrieves public biological knowledge | `pathway_context`,<br>`gene_disease_associations`,<br>`search_papers` | g:Profiler, Reactome, Open Targets, GWAS Catalog, Europe PMC |
| Internal analysis | Performs local computation and figure generation | `analyze_deseq2_counts`,<br>`run_expression_pca`,<br>`generate_figure` | DESeq2, pandas, scikit learn, matplotlib, local R scripts |
| Workflow management | Manages sessions, retained materials and reports | `create_session`, `get_artifact`,<br>`create_session_report` | Local workspace store, report templates |
| Hybrid orchestration | Combines local analysis with external evidence retrieval | `analyze_rnaseq_result`,<br>`enrich_gene_set`,<br>`investigate_gene_set` | Local RNA sequencing parser, clusterProfiler, ReactomePA, DOSE, g:Profiler, Open Targets, Europe PMC |

## References

[1] Chen, H. et al. Scaling and quantization of large-scale foundation model enables resource-efficient predictions in network biology. Nature Computational Science 6, 450–463 (2026). 10.1038/s43588-026-00972-4

[2] Qu, K. et al. Integrative genomic analysis by interoperation of bioinformatics tools in GenomeSpace. Nature Methods 13, 245–247 (2016). 10.1038/nmeth.3732

[3] Reich, M. et al. GenePattern 2.0. Nature Genetics 38, 500–501 (2006). 10.1038/ng0506-500

[4] Afgan, E. et al. The Galaxy platform for accessible, reproducible and collaborative biomedical analyses: 2018 update. Nucleic Acids Research 46, W537–W544 (2018). 10.1093/nar/gky379

[5] Hull, D. et al. Taverna: a tool for building and running workflows of services. Nucleic Acids Research 34, W729–W732 (2006). 10.1093/nar/gkl320

[6] Reich, M. et al. The GenePattern Notebook Environment. Cell Systems 5, 149–151.e1 (2017). 10.1016/j.cels.2017.07.003

[7] Torre, D., Lachmann, A. & Ma’ayan, A. BioJupies: automated generation of interactive notebooks for RNA-seq data analysis in the cloud. Cell Systems 7, 556–561.e3 (2018). 10.1016/j.cels.2018.10.007

[8] Yao, S., et al. ReAct: synergizing reasoning and acting in language models. International Conference on Learning Representations (2023). https://openreview.net/forum?id=WE_vluYUL-X

[9] Schick, T., et al. Toolformer: language models can teach themselves to use tools. Advances in Neural Information Processing Systems 36 (2023). https://proceedings.neurips.cc/paper_files/paper/2023/hash/d842425e4bf79ba039352da0f658a906-Abstract-Conference.html

[10] Jin, Q. et al. GeneGPT: augmenting large language models with domain tools for improved access to biomedical information. Bioinformatics 40, btae075 (2024). 10.1093/bioinformatics/btae075

[11] Nasri, F. et al. Deterministic access to global viral sequence data enables robust agentic scientific discovery. arXiv preprint arXiv:2606.06749 (2026). 10.48550/arXiv.2606.06749

[12] Huang, K. et al. Autonomous biomedical research with an artificial intelligence agent. Science 393, eadz4351 (2026). 10.1126/science.adz4351

[13] Chen, Z. et al. An agentic artificially intelligent X-ray scientist. Nature Machine Intelligence 8, 1075–1086 (2026). 10.1038/s42256-026-01261-5

[14] Ding, K. et al. SciToolAgent: a knowledge-graph-driven scientific agent for multitool integration. Nature Computational Science 5, 962–972 (2025). 10.1038/s43588-025-00849-y

[15] Shao, E. et al. SciSciGPT: advancing human–AI collaboration in the science of science. Nature Computational Science 6, 301–315 (2026). 10.1038/s43588-025-00906-6

[16] Ghareeb, A. E. et al. A multi-agent system for automating scientific discovery. Nature 655, 497–505 (2026). 10.1038/s41586-026-10652-y

[17] Gottweis, J. et al. Accelerating scientific discovery with Co-Scientist. Nature 655, 487–496 (2026). 10.1038/s41586-026-10644-y

[18] Lu, C. et al. Towards end-to-end automation of AI research. Nature 651, 914–919 (2026). 10.1038/s41586-026-10265-5

[19] Model Context Protocol. Model Context Protocol documentation. https://modelcontextprotocol.io/ (accessed 16 September 2026).

[20] Mitchener, L. et al. BixBench: a comprehensive benchmark for LLM-based agents in computational biology. arXiv preprint arXiv:2503.00096 (2025). 10.48550/arXiv.2503.00096

[21] OpenCode [software]. GitHub, https://github.com/anomalyco/opencode (accessed 17 September 2026).

[22] Model Context Protocol. Specification: transports (2025). https://modelcontextprotocol.io/specification/2025-06-18/basic/transports

[23] Love, M. I., Huber, W. & Anders, S. Moderated estimation of fold change and dispersion for RNA-seq data with DESeq2. Genome Biology 15, 550 (2014). 10.1186/s13059-014-0550-8

[24] Moraes, A. & Ferreira, P. GLESKit supplementary data and evaluation records. Zenodo (2026). 10.5281/zenodo.23592806

